# High-Resolution Imaging of the Rhizosphere Using Resin-Embedded Cross-Sectional Polishing and Scanning Electron Microscopy for Biological Samples (bioCP-SEM)

**DOI:** 10.1101/2025.11.27.690849

**Authors:** Kiminori Toyooka, Yuko Saito, Yumi Goto, Mayuko Sato

## Abstract

Understanding microbial colonization in plant-associated environments requires high-resolution imaging of intact plant-microbe interfaces under natural soil conditions. However, conventional electron microscopy is limited by sample distortion and poor compatibility with heterogeneous substrates. Here, we present bioCP-SEM, a cross-sectional polishing scanning electron microscopy technique optimized for biological specimens inhabiting unsterilized soil. This method integrates resin embedding, diamond band saw sectioning, and mechanical polishing to enable high-quality imaging across large, complex samples. Using bioCP-SEM, we visualized bacterial clusters in a model plant root, and multilayered biofilm-like structures and fungal structures, including arbuscule-like forms, in a wild weed species. We further applied this method to intact moss collected from an urban environment, capturing spatially resolved interactions among biofilms, spores, protozoa, and annelids. Notably, microbial cells and subcellular structures, such as cilia, were preserved in situ. These results demonstrate the versatility of bioCP-SEM for mapping microbial architecture across diverse biological and environmental contexts.

## Main

The rhizosphere is a biologically active zone where plants interact with diverse microbial consortia, including biofilm-forming bacteria and symbiotic mycorrhizal fungi^1–4^. Biofilms, composed of microorganisms embedded in extracellular polymeric substances (EPS), provide structural integrity, retain nutrients, and protect soil microbiota from environmental stressors^5–6^. Additionally, ectomycorrhizal (ECM) and arbuscular mycorrhizal (AM) fungi form symbiotic interfaces with plant roots that facilitate mineral acquisition and immune modulation^7–9^. Despite their ecological importance, direct ultrastructural visualization of biofilms and mycorrhizae in intact soil–root systems remains largely inaccessible using conventional methods^10–12^. Recently, ultramicrotome sectioning and transmission electron microscopy (TEM) have been applied to visualize soil ultrastructures at the nanometer scale, revealing close spatial associations between microbes, minerals, and organic matter^13^. However, such approaches remain limited by field of view and sample preparation constraints. Furthermore, when using an ultramicrotome to prepare ultrathin sections from samples containing minerals, many problems arise, such as hard minerals scratching diamond knives and preventing the acquisition of smooth surface sections.

To address this methodological gap, we sought to establish a cross-sectional polishing scanning electron microscopy (SEM) workflow tailored for large, resin-embedded biological–mineral composites. Building upon advances in cross-sectional polishing used in materials science, we adapted diamond band-saw sectioning and automated mechanical polishing for biological samples to enable the preparation of smooth, wide-area surfaces suitable for high-resolution SEM. This workflow—comprising fixation, resin embedding, sectioning, polishing, staining, and field emission SEM (FE-SEM) observation—is schematically summarized in Fig. 1 and provides a basis for analyzing rhizosphere samples containing both mineral and organic materials.

**Figure 1.**
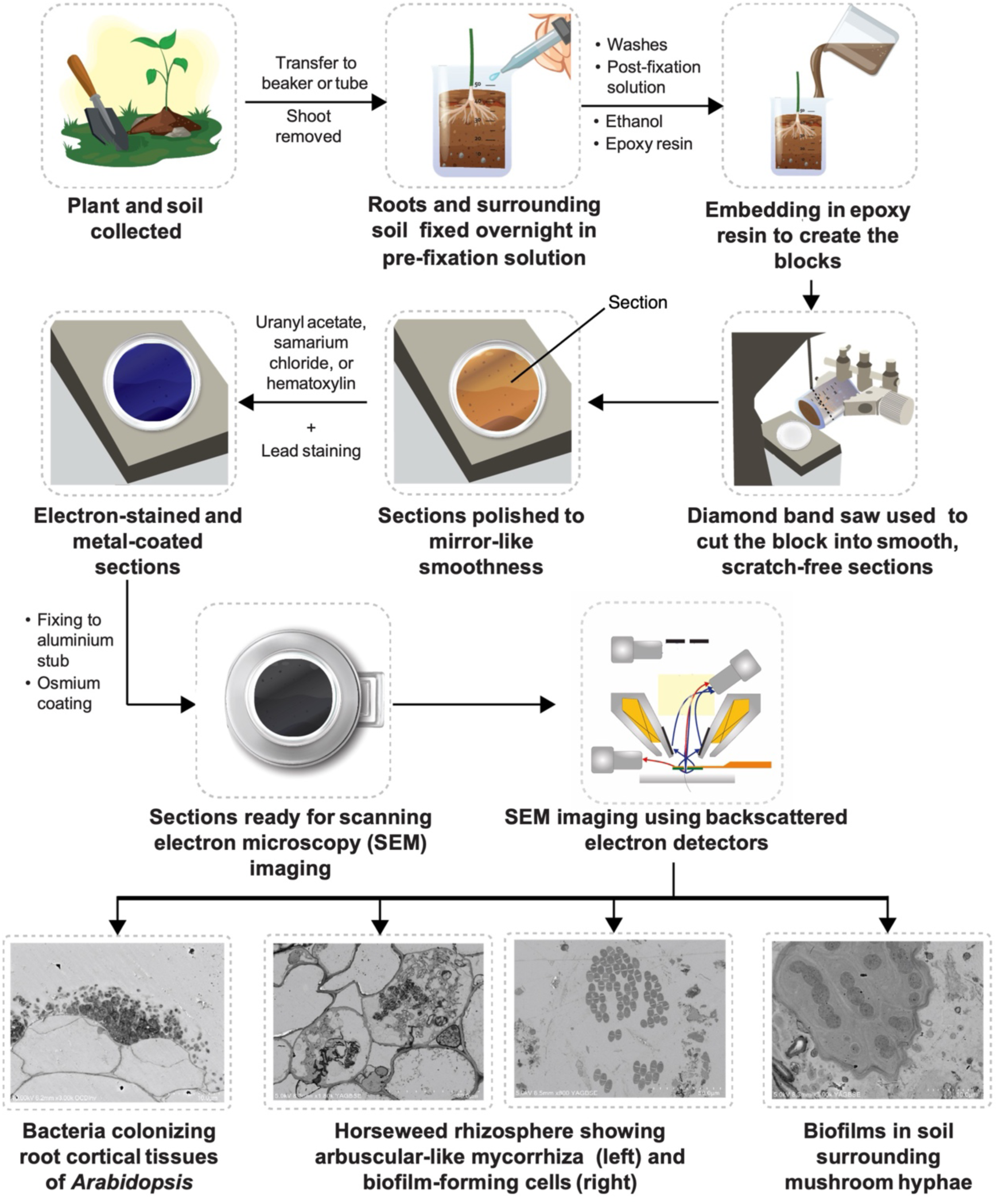
Graphical Abstract. Schematic overview of the bioCP-SEM method showing field collection of plant–soil samples, resin embedding, cross-sectional polishing, and high-resolution SEM imaging, revealing plant–microbe structures.

Recent studies have highlighted that plant tissues grown in natural, non-sterile soils harbor diverse microbial assemblages, including bacteria and fungi that inhabit root surfaces and intercellular regions^1–2^. However, visualizing these associations in situ remains technically challenging, particularly when attempting to preserve the spatial context of microbial consortia within heterogeneous plant–soil environments. A method capable of resolving microbial architecture across both model and non-model plant species—while maintaining intact interfaces between roots, tissues, and surrounding soil components—would substantially advance our understanding of plant–microbe interactions under natural conditions.

The capacity to visualize both external and internal fungal symbioses alongside biofilm development in situ offers a powerful new perspective on spatial microbial organization. BioCP-SEM enables comparative ultrastructural analyses of rhizosphere microbial architecture across species and environmental contexts, bridging the gap between ecological function and spatial form.

Accordingly, the aim of this study was to develop and validate a cross-sectional polishing SEM method that enables spatially preserved, high-resolution visualization of microbial structures within intact soil–root systems. Specifically, we sought to (i) establish a workflow for preparing large resin-embedded plant–soil composites, (ii) evaluate its applicability across both model and non-model plant species grown in natural soils, and (iii) provide a methodological foundation for comparative ultrastructural analyses of biofilms, fungal symbioses, and other microbe–tissue interfaces in heterogeneous rhizosphere environments.

In this study, organismal identities inferred from SEM were based solely on the morphology and context observed on backscattered electron (BSE) images. We therefore employ descriptive qualifiers, such as “-like,” “putative,” or “presumed” (e.g., “protozoa-like,” “arbuscule-like,” “annelid-like”), and avoid species- or genus-level claims without molecular or lineage-specific markers. This cautious style follows precedents in environmental electron microscopy studies of difficult-to-classify microbiota and is appropriate for heterogeneous soil composites^14^.

## Results

### Efficient preparation of large, resin-embedded rhizosphere samples

The bioCP-SEM method enabled well-preserved preparation of cross sections from complex biological–mineral composites. To preserve native spatial structure, we collected entire root–soil systems either by extracting root zones from pots or by boring soil blocks containing wild plants. These samples were placed in 5-, 14- or 50-mL polypropylene tubes or in 100 200-mL polypropylene beakers with a drainage hole at the bottom. Pre-fixation was carried out using a solution containing 4% formaldehyde and 2% glutaraldehyde. The solution was gently added from the top, similar to watering plants, to allow gradual infiltration without disturbing the sample structure (Fig. 2a). After pre-fixation, samples were degassed to improve penetration of fixatives and buffer into the soil matrix. Degassing was performed by reducing gage pressure to approximately - 0.08∼-0.1 MPa for several short cycles, followed by a final cycle in which samples were held under vacuum for 30 min before slowly returning to atmospheric pressure. Following an overnight fixation at 4 °C, solutions—including washes, post-fixation solution containing 1% osmium tetroxide, graded ethanol, and embedding resin—were successively exchanged through the bottom hole while fresh solution was added from the top. This gravity-assisted fluid exchange ensured complete infiltration of a low-viscosity epoxy resin (Quetol 651). After the transition to 100% resin, a second degassing step was performed using the same procedure (several short cycles to 0.08 MPa followed by 30 min under vacuum) to remove residual trapped air and facilitate full polymer infiltration throughout the sample. This combination of gravity-driven exchange and staged degassing minimized the formation of voids and yielded large, resin-embedded blocks suitable for polishing and imaging.

**Figure 2.**
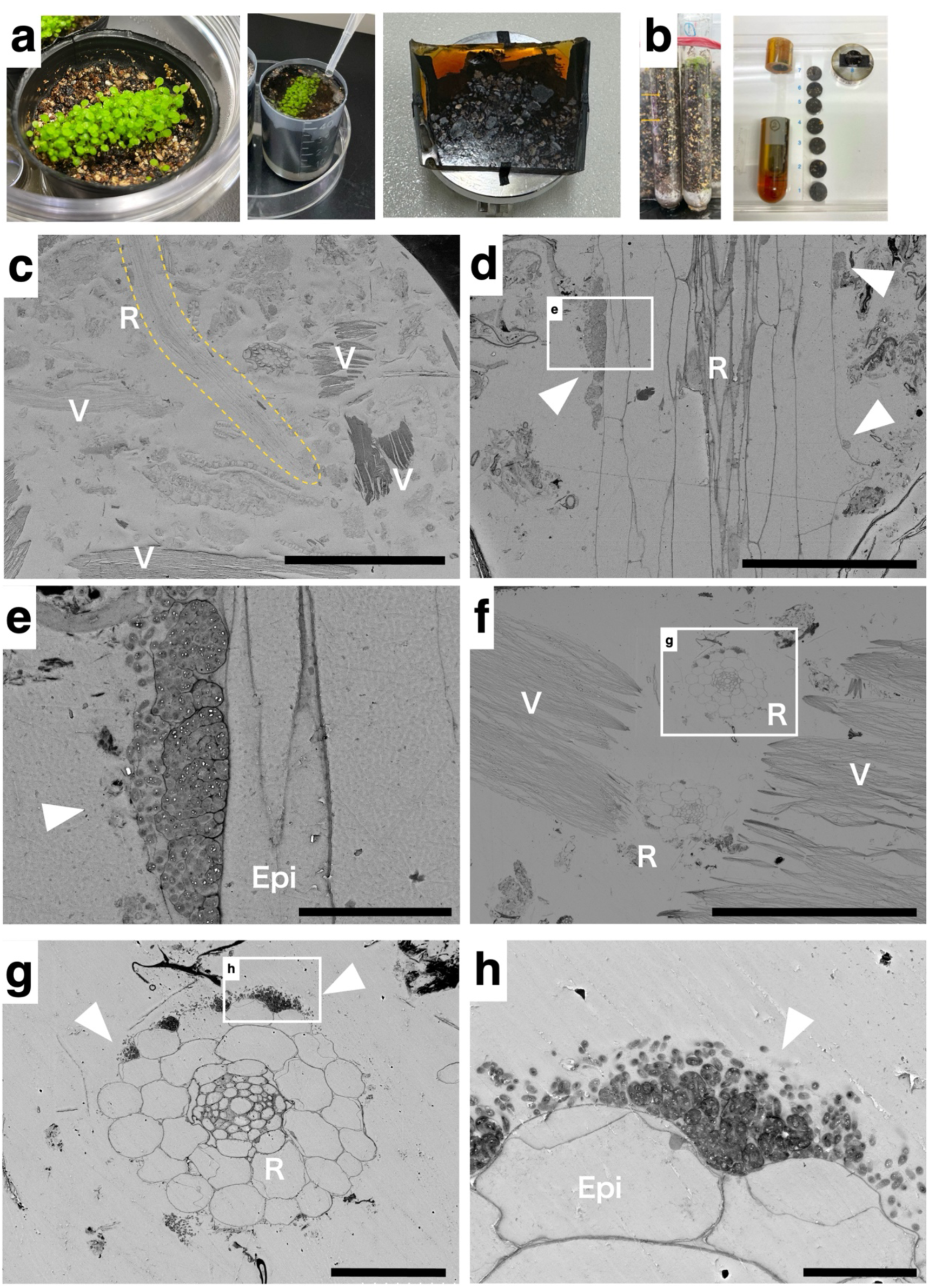
Rhizosphere imaging of Arabidopsis roots. (a, b) Preparation of resin-embedded soil samples containing Arabidopsis roots. (a) Arabidopsis seedlings were grown in polypropylene cups filled with non-sterile soil (a mixture of vermiculite and peat moss) and fixed by gently adding pre-fixation solution from the top. The resin-embedded blocks were longitudinally sectioned, and the cross-sections were polished, stained, and observed (c–e). (b) Arabidopsis seedlings were also cultivated in conical tubes containing non-sterile soil. After resin embedding, the blocks were cut into 2-mm slices, and the uppermost section was selected for observation (f–h). (c) Low-magnification view showing root (R, outlined in yellow) surrounded by vermiculite particles (V). (d, e) Longitudinal section of the root maturation zone. Arrowheads indicate microbial aggregates associated with the root surface. (e) Higher magnification of the boxed area in (d), showing root epidermis (Epi) and attached bacterial cells (arrowhead). (f, g) Broader views of the root–vermiculite interface. (f) Root (R) in contact with vermiculite (V); boxed area corresponds to (g). (g) Cross-sectional image of root tissues; arrowheads indicate bacterial clusters near the epidermal surface. (h) Higher magnification of the boxed area in (g), showing bacterial aggregation (arrowhead) attached to epidermal cells (Epi). Scale bars: (c) 500 µm, (d, g) 50 µm, (e, h) 10 µm, (f) 300 µm.

To accommodate the hardness and heterogeneity of resin-embedded soil samples, we used a diamond band saw, commonly applied for cutting mineralized biological specimens such as bone and teeth. The obtained sections (3–5 mm thick, across cm-scale blocks) exhibited surface scratches, which made direct imaging of ultrastructural details impossible without subsequent polishing or milling (Extended Data Fig. 1a, b). To achieve a mirror-like cross-sectional surface, we optimized a mechanical polishing machine using two-stage abrasive paper and a polishing protocol (Extended Data Fig. 1c, d). If polishing was insufficient, we used diamond slurry or an ion milling machine to perform final polishing (Extended Data Fig. 1e–h). This combination of industrial-grade cutting and polishing preserved the fragile root and microbial structures and enabled consistent imaging across a wide range of field-collected samples (Fig. 1).

To enhance contrast and minimize electron charging during SEM imaging, sections were subjected to electron staining and conductive coating. Optimal contrast was achieved using sequential uranyl acetate and lead staining, while non-radioactive alternatives—such as samarium chloride^15^ or Meyer’s hematoxylin^16^—provided comparable results. Dried sections were fixed to aluminum stubs with carbon adhesive and coated with osmium, allowing stable imaging without charging artifacts (Extended Data Fig. 2). Full protocols and comparative results are detailed in the Supplementary Information.

### High-resolution imaging of rhizosphere ultrastructure

SEM imaging was performed using BSE detectors, where regions rich in heavy elements appear bright and those composed of lighter elements appear dark^17^. This process produces an inverted image contrast relative to that of a conventional TEM image. This inversion may result in an unfamiliar appearance for those who are accustomed to TEM micrographs. To facilitate interpretation, grayscale inversion was applied using the SEM operation software option to produce positive-contrast images resembling TEM views. Alternatively, BSE images were inverted to black and white using image processing software (ImageJ, Fiji). Among BSE detector types, the Yttrium Aluminum Garnet (YAG)-BSE detector (or its successor, the out-column crystal type BSE detector, OCD-BSE) provided higher responsivity and contrast than the conventional in-lens BSE detector (often referred to as a low-angle BSE detector in earlier Hitachi models) when imaging samples with mixed organic and mineral content, such as vermiculites and root structures. In addition, despite reduced resolution, bioCP-SEM observation was possible with the latest general-purpose and tabletop SEMs equipped with BSE detectors.

Using these optimized conditions, we visualized the rhizosphere of *Arabidopsis thaliana* grown in unsterilized soil (Fig. 2a–c). Vertical and horizontal section images revealed bacterial clusters and structured biofilm-like layers associated with the root surface (Fig. 2d–h).

In wild plants [e.g., Horseweed (*Erigeron* sp.)] collected from the landscaped area shown in Fig. 3a, extensive microbial colonization was observed around the lateral root, including clusters of cocci-shaped microorganisms surrounding the hypha-like filaments and organic matter of ectomycorrhizal fungi (Fig. 3b–d), and arbuscular-like structures developing within the root cortex cells (Fig. 3e–g). Furthermore, compared with the other methods, bioCP-SEM provided the highest resolution capable of identifying ribosomes associated with the endoplasmic reticulum (Fig. 3g). In the aboveground portion of the main root of the same plant, deformed plant tissue—such as a gall—was observed (Fig. 3h–i). Proliferating microorganisms were identified on the epidermal cell surface, and the acquired images suggested that microorganisms loaded onto plant cell wall-derived cargo were spreading into the soil (Fig. 3j–k). Among these, we also observed an encapsulated microbial structure morphologically reminiscent of a spore or a testate amoeba (Fig. 3l), although its precise identity could not be determined from morphology alone. The rhizosphere revealed a diverse array of unknown microorganisms, including cells forming biofilm-like matrices (Fig. 3m–p). The observation of the internal structure of biofilms using electron microscopy has proven to be a challenging procedure.

**Figure 3.**
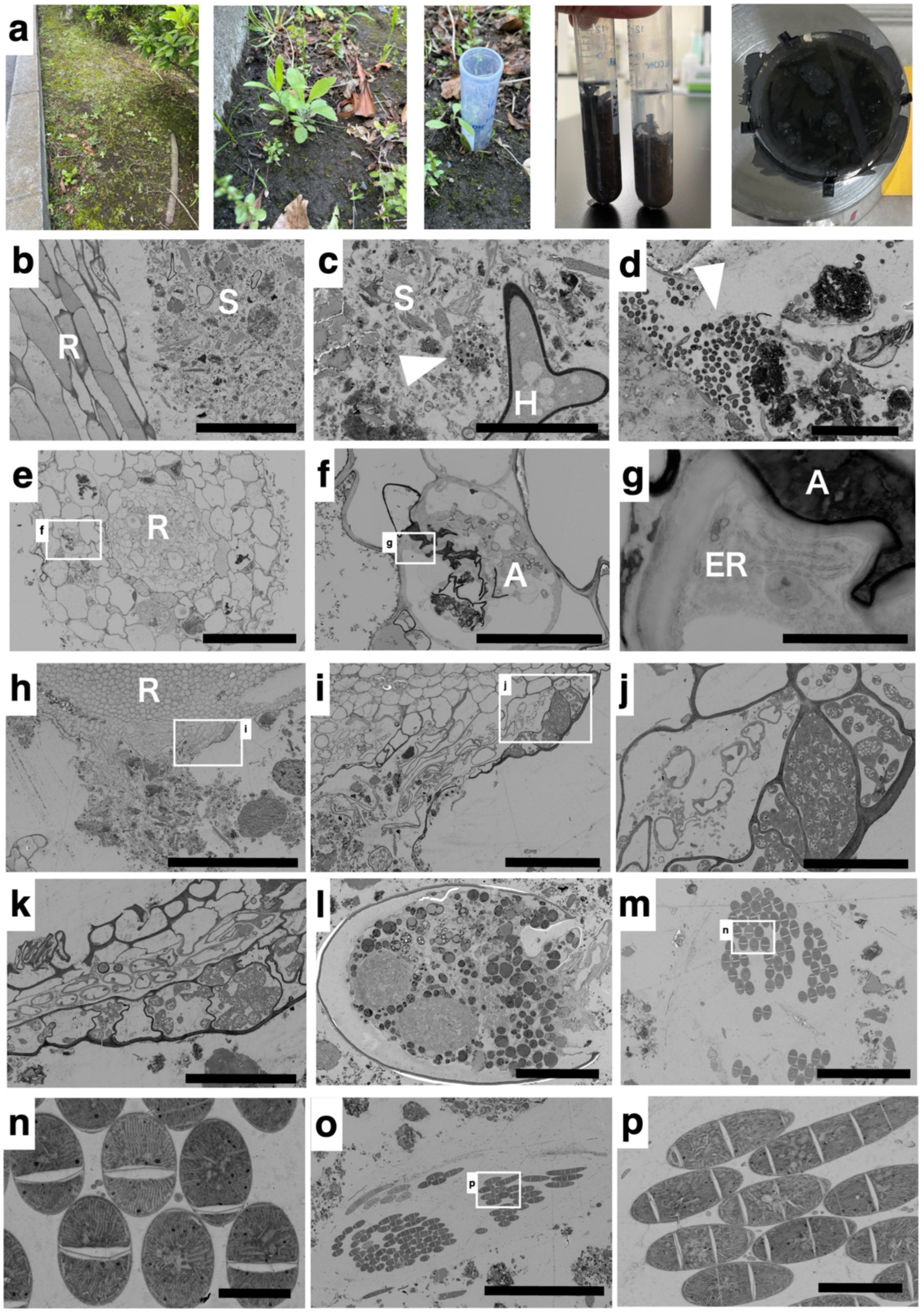
Ultrastructural views of giant horseweed. (a) Giant horseweed (*Erigeron* sp., species not identified) was collected from a landscaped area using a bottomless conical tube and then embedded in resin for bioCP-SEM observation. (b) Longitudinal section of root (R) and surrounding soil (S). (c) Soil and hypha (H) associated with bacterial cells (arrowhead). (d) Higher magnification showing clusters of bacteria (arrowhead). (e–g) Lateral root (R) infected with arbuscular mycorrhizal (AM) fungi. (f) Enlarged view of boxed region in (e), showing fungal arbuscule (A) within a host cell. (g) Higher magnification of the boxed area in (f), showing arbuscule (A) closely associated with the endoplasmic reticulum (ER). (h–k) Gall-like structure observed in the above-ground portion of the main root. (i) Enlarged view of the boxed region in (h), showing hypertrophied host cells colonized by microorganisms. (j, k) Further magnified views of the boxed region in (i). (l) An encapsulated microbial structure morphologically reminiscent of a spore or a testate amoeba, although its precise identity remains unknown. (m–p) Unknown microorganisms, including cells forming biofilm-like matrices. (n) Higher magnification of the boxed area in (m), showing biofilm-forming cells with regular ultrastructure. (p) Higher magnification of the boxed area in (o), showing elongated biofilm-forming cells. Arrowheads indicate aggregation of bacteria. Abbreviations: A, Arbuscule; AM, Arbuscular mycorrhiza; Epi, Epidermis; ER, Endoplasmic reticulum; H, Hypha; R, Root; S, Soil. Scale bars: (h) 500 µm; (b, e, i, o, r) 100 µm; (k, p) 50 µm; (j) 30 µm; (c, f, l) 20 µm; (d, n, s) 10 µm; (m, q) 5 µm; (g) 2 µm.

As shown in Fig. 4a, various microorganisms in biofilms were observed in the soil where mushrooms naturally grow. Algae and bacteria biofilms, basidiomycete, and aggregates of spores and bacteria were observed around the leafy gametophyte of moss (species not identified) and in the soil (Fig. 4b–g). At the base of cherry sprouts (Fig. 4h, i), endophytic fungi (Fig. 4k) and microbial aggregates associated with the root surface were detected (Fig 4l, m). These findings demonstrate that bioCP-SEM is a versatile method capable of visualizing not only soil-associated microorganisms but also lichens growing on rocks^21^ and outdoor moss, including soil dust.

**Figure 4.**
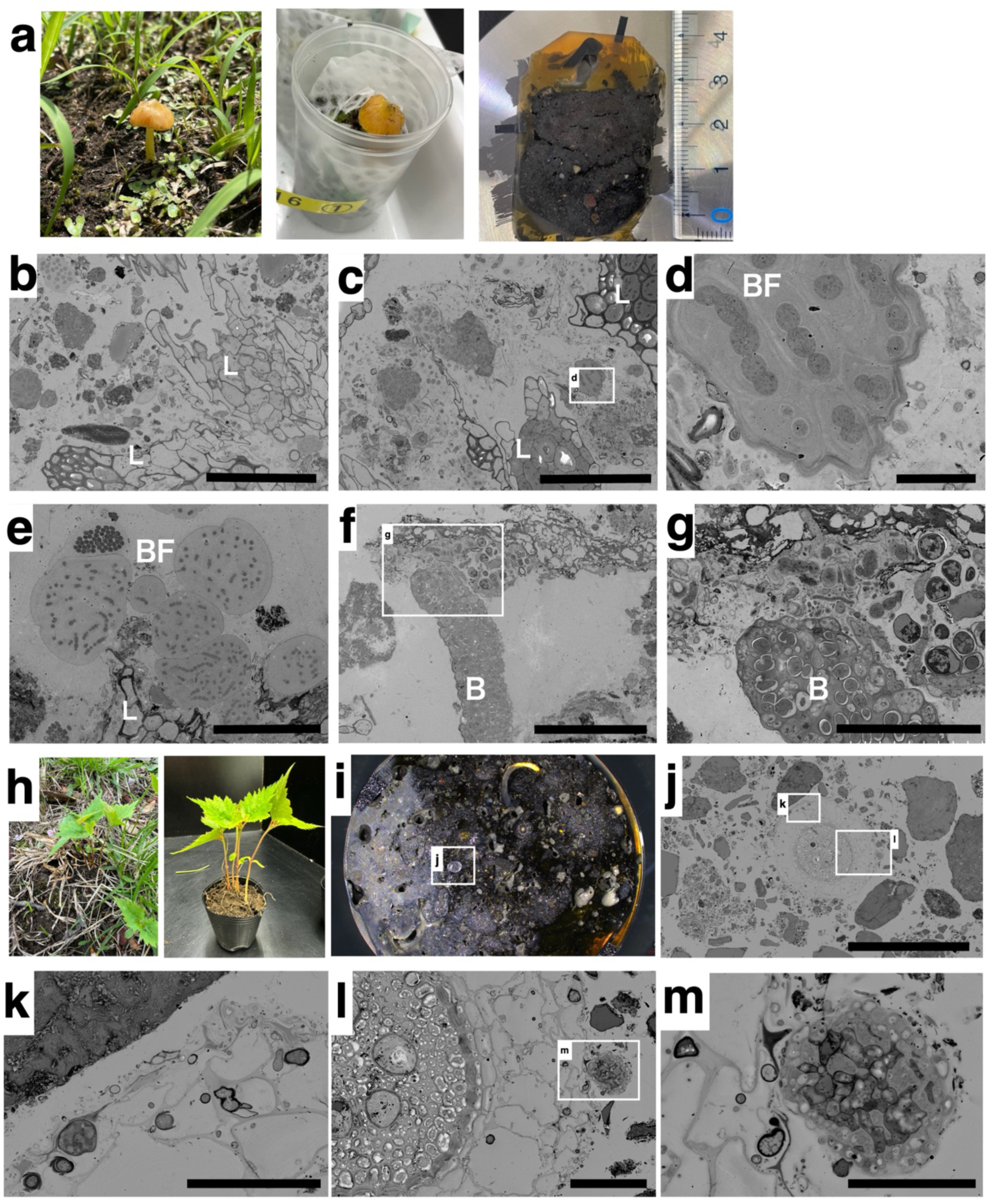
Imaging of the rhizosphere of a mushroom with underlying mosses and a cherry blossom seedling. (a–d) Fungal and bacterial associations. (a) Mushroom and underlying mosses (species not identified) collected from a landscaped area and embedded for bioCP-SEM observation. (b) Polished section of moss leaf (L) tissue showing associated microbes. (c) Cross-section of moss leaves (L) and soil, showing microbial colonization. (d) Biofilm-like (BF) structures formed by endophytic bacteria within the moss. (e–g) Balloon-like microbial inclusions and biofilm structures. (e) Microbial aggregates forming a layered BF near the moss tissue (L). (f) Enlarged view of microbial clusters (B) attached to the tissue surface; boxed area corresponds to (g). (g) Higher magnification showing balloon-like microbial inclusions and close microbial associations. (h–m) Rhizosphere and bud tissues of cherry blossoms (species not identified). (h) Cherry sprouts collected from a landscaped area were replanted into a pod. (i) Resin-embedded soil and root sample block. (j) Low-magnification view of the rhizosphere section; boxed areas correspond to (k) and (l). (k) Root-associated microbial communities. (l) Enlarged view of rhizosphere with dense microbial colonization; boxed area corresponds to (m). (m) Higher magnification showing clustered microbial cells forming biofilm-like structures. Abbreviations: B, Bacteria; BF, Biofilm; L, Leaves of mosses. Scale bars: (j) 400 µm; (b, c, e, f) 100 µm; (g, l) 50 µm; (k) 40 µm; (m) 20 µm; (d) 10 µm.

Fig. 5 shows bioCP-SEM images of a colony of moss (species not identified) growing on a roadside grate. The moss colony was fixed, embedded in its entirety, polished, and observed (Fig. 5a). Numerous biofilms, bacterial clusters, and spores were observed around the moss gametophyte leaves (Fig. 5b–e). Further observations revealed the presence of protozoa-like cells and a small annelid-like organism (Fig. 6). Cross-sectional imaging captured numerous intracellular vesicles (Fig. 6a) and intricate membrane structures in protozoa-like cells (Fig. 6b), as well as ciliated epithelial tissues densely covered with motile cilia and microvilli-like protrusions in the annelid-like organism (Fig. 6c–d). These features are consistent with a ciliated epithelial organ (e.g., nephridial/excretory duct), rather than a digestive tract. Surrounding soil matrices contained diverse microorganisms, including bacteria and fungi (Fig. 6e–g). These observations demonstrate that bioCP-SEM enables high-resolution visualization of multicellular and microbial structures in complex soil environments.

**Figure 5.**
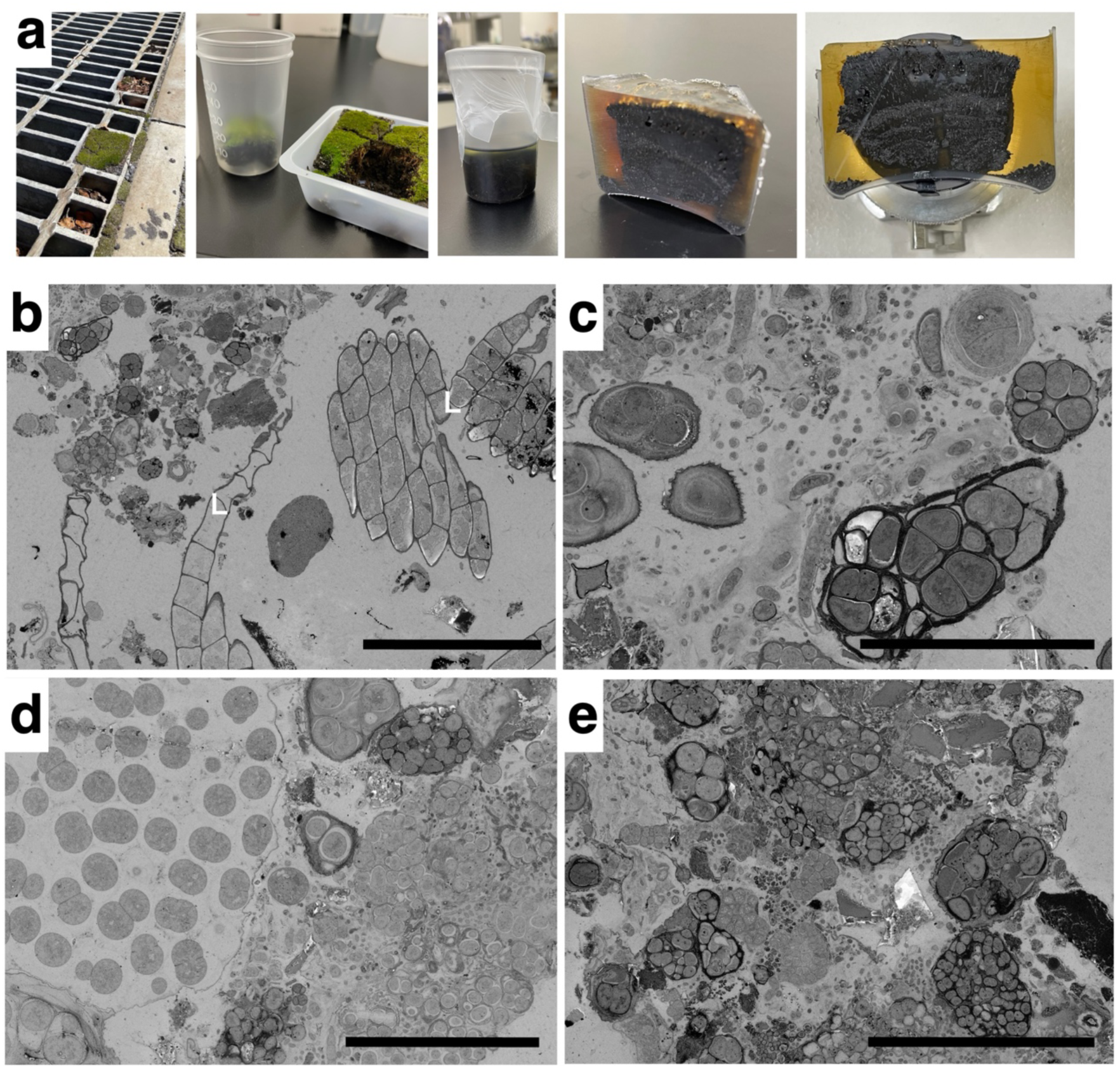
Sample preparation of moss growing and biofilm-associated organisms in moss-covered soil. (a) Moss colony growing in the crevices of a roadside metal grating were sampled (species not identified). Collected moss–soil blocks were immersed in fixative solution in polypropylene cups. After fixation and dehydration, the samples were embedded in epoxy resin. Cross-sections of resin-embedded moss–soil blocks were prepared using the bioCP-SEM method. (b–e) Cross-sectional views of moss leaves (L) and associated microbial communities. Around the basal area of leafy shoots were observed. (b) Moss leaf tissues (L) surrounded by soil particles; numerous microbial aggregates are present in the extracellular spaces. (c–e) Higher magnification views showing diverse microbial cells and biofilm-like structures closely associated with the moss tissues. Numerous small, round particles observed between the moss leaves correspond to diverse microorganisms. Scale bars: (b) 100 µm; (c, d) 30 µm; (e) 40 µm.

**Figure 6.**
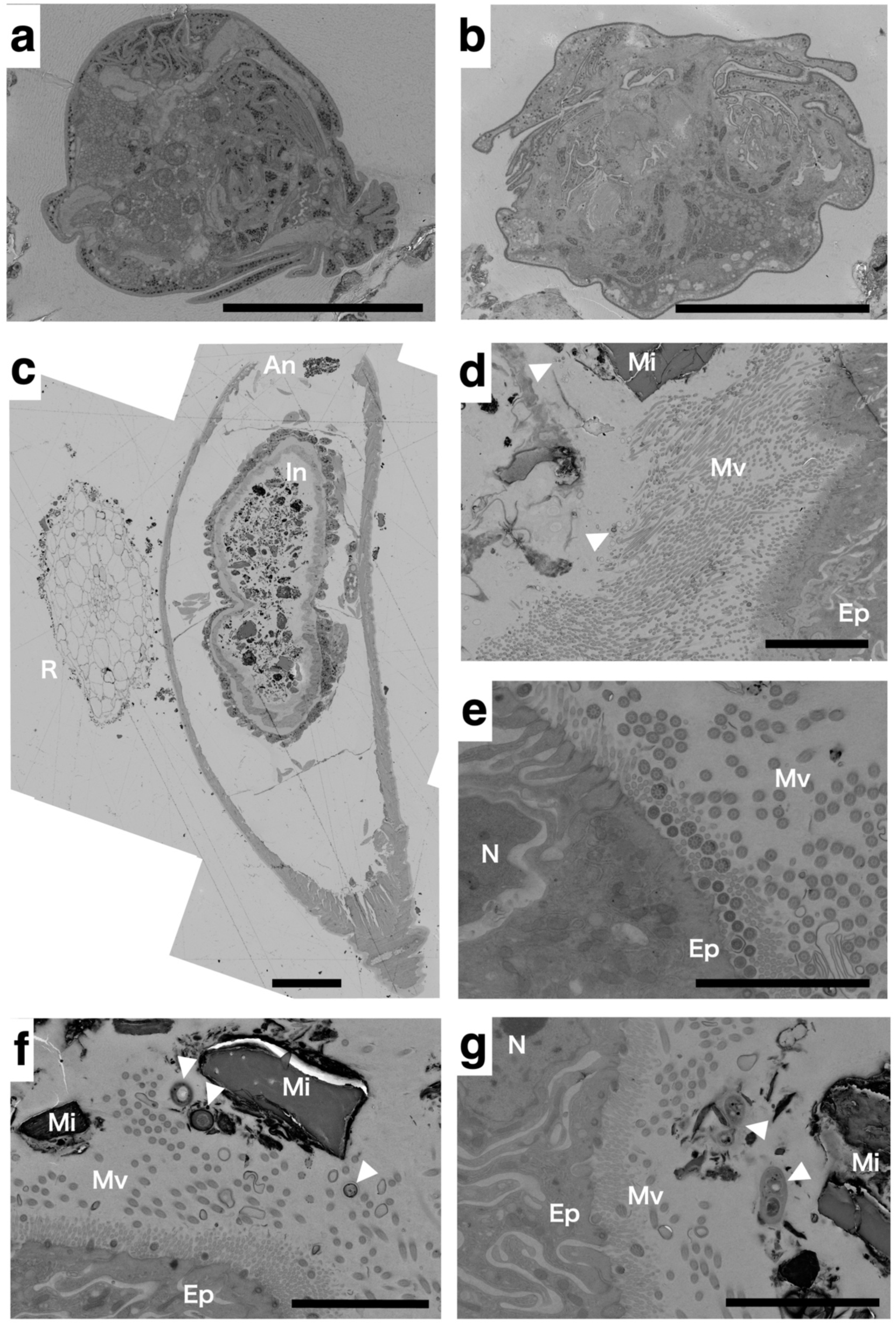
Protozoa and small annelid in natural soil. (a, b) Ultrastructure of protozoa observed in soil samples, showing intracellular vesicles and membrane structures. (c) Whole cross-sectional view of an annelid (An) intestine (In) adjacent to a plant root (R). (d–g) Ultrastructures of intestinal villi and microvilli (Mv) of the annelid. (d) Overview of intestinal wall and attached bacteria (arrowheads). (e) Higher magnification of epidermal cells (Ep) with nuclei (N) and aligned microvilli (Mv). (f) Microbial cells (arrowheads) and mineral particles (Mi) closely associated with the intestinal surface. (g) Microvilli (Mv) in contact with mineral particles (Mi) and bacteria (arrowheads). Arrowheads indicate bacteria. Abbreviations: Ep, Epidermal cell; An, Annelid; In, Intestine; Mi, Mineral; Mv, Microvilli; N, Nucleus; R, Root. Scale bars: (a, b) 30 µm; (c) 100 µm; (d) 20 µm; (e) 4 µm; (f, g) 5 µm.

## Discussion

The bioCP-SEM method provides a robust platform for imaging plant–soil interfaces in situ. By adapting industrial-grade cutting and polishing tools for biological SEM, this technique overcomes key limitations of ultramicrotomy and FIB-SEM^19–22^, enabling cross-sectional imaging across large and heterogeneous samples. The resulting preservation of spatial context, microbial diversity, and root architecture supports integrative analyses of rhizosphere structure and function.

From a technical standpoint, sample fixation and resin infiltration are critical for ensuring adequate preservation of fragile microbial and plant tissues within non-sterile soils. The gradual infiltration of resin into soil aggregates minimizes collapse and shrinkage, thereby maintaining spatial relationships between roots and surrounding microorganisms. Furthermore, the introduction of a diamond band saw—originally developed for hard tissue and material science applications—represents a novel adaptation to soil-sectioning for biological SEM. This approach enables the reproducible production of thick sections (2–5 mm), which retain sufficient structural integrity for subsequent polishing and staining, while providing a broad field of view for microbial consortia analysis (Fig. 2).

Recent advances in BSE detection, particularly the development of high-sensitivity detectors, have significantly improved imaging resolution. These improvements now allow for subcellular features to be resolved on cross-polished surfaces, achieving image quality comparable to that of TEM while retaining a broader field of view. This enhancement further expands the applicability of the bioCP-SEM method to ultrastructural studies.

However, with the expansion of the observable surface area, the consumption of heavy metal stains, such as uranyl acetate, substantially increases. To address this limitation, we used Mayer’s hematoxylin as a non-radioactive alternative contrast agent^16^. It provided sufficient electron density for BSE-based imaging of biological structures, making it a viable substitute for large-scale rhizosphere imaging.

In Arabidopsis roots grown in unautoclaved soil, previous metagenomic analyses and imaging of the root zone have demonstrated dense bacterial colonization and aggregates in the intercellular spaces between epidermal cells^23–25^. Our bioCP-SEM analysis likewise revealed clusters of bacteria on the Arabidopsis root surface and within the intercellular spaces, consistent with these earlier observations. This agreement across sequencing- and imaging-based approaches supports the view that Arabidopsis roots in natural soils are extensively colonized by diverse bacterial communities (Fig. 2) and highlights the potential of bioCP-SEM to validate and extend findings from fluorescence-based microbiome studies^23–25^.

We observed an abundance of microbial morphotypes—rhizobia, mycorrhizal fungi, and dense biofilms—within intact soil–root systems. Notably, the workflow also revealed microbial communities and structural phenomena that have been difficult to visualize with conventional methods (Fig. 3). These findings underscore the ecological complexity of the rhizosphere and demonstrate the value of ultrastructural imaging in uncovering novel plant–microbe interactions. These structures further illustrate how thick-section polishing can preserve microbial consortia without disrupting their native architecture. Future experiments could leverage this thickness to systematically compare microbial community organization across plant species, soil types, or environmental conditions.

It is estimated that 90%–99% of soil microorganisms remain uncultured and taxonomically uncharacterized, highlighting the vast unknown microbial diversity in terrestrial ecosystems^26,27^. Originally reported by Torsvik et al. ^26^ and further emphasized by Daniel^27^, this gap reflects the limitations of culture-based approaches and underscores the need for in situ visualization techniques. BioCP-SEM provides a direct means to access this hidden microbiota, potentially enabling morphological identification and spatial mapping of yet-uncultured organisms (Figs. 3–5).

In the resin-embedded soil samples, a small annelid-like organism displayed epithelial tissues densely covered with motile cilia and microvilli-like protrusions (Fig. 6). The presence of 9+2 axonemal structures indicates active motility, suggesting that these cilia serve a transport or pumping function rather than the absorptive role typically associated with intestinal microvilli. In many oligochaete groups, ciliated epithelia with mixed apical specializations are known to occur in excretory or filtrative organs. The structure observed here is therefore more consistent with a ciliated duct, such as excretory channel, than with digestive tissue. Definitive identification, however, will require molecular or developmental characterization of the organism.

In the future, the development of curated image libraries of ultrastructural signatures will further accelerate correlative imaging workflows^28^ and support emerging frameworks for artificial-intelligence–driven microbial detection and taxonomic classification^29^. By linking structural phenotypes—such as organelle-mediated transport pathways^30^, biofilm organization^3^ and microbe–host interfaces—with microbial diversity, bioCP-SEM will support high-throughput, spatially resolved analyses of rhizosphere ecology.

Despite these advantages, several methodological limitations should be acknowledged. Resin infiltration remains a major constraint when working with dense or water-retentive soils, where fixatives and epoxy resins penetrate unevenly. In preliminary trials using large rhizosphere samples embedded in 500-mL beakers, incomplete resin polymerization occurred, resulting in fragile blocks that were difficult to section or polish. These challenges suggest that optimized dehydration strategies, pre-conditioning of soil aggregates, or staged infiltration protocols may be required to ensure complete polymerization in large or compacted samples. Moreover, organismal identification in heterogeneous soil composites is inherently limited by morphology-based inference; integrating molecular markers, isotopic labeling, or single-cell genomic approaches will be essential for more definitive classification. Addressing these methodological constraints will broaden the applicability of bioCP-SEM and enable more quantitative, scalable analyses of plant–soil–microbe associations across diverse environmental contexts.

Finally, the bioCP-SEM workflow offers substantial practical advantages. Because it relies on standard epoxy resins, automated mechanical polishing, and commonly available SEM instrumentation, the method can be implemented in most electron microscopy facilities without requiring specialized equipment or advanced ultramicrotomy skills. The elimination of ultrathin sectioning—one of the most technically demanding steps in conventional ultrastructural analysis—greatly lowers the barrier to adoption. This accessibility extends the utility of the approach beyond plant roots to animal tissues, complex biofilms, and diverse soil microbial consortia. Consequently, bioCP-SEM provides a versatile, widely usable platform for high-resolution ultrastructural studies across biological and environmental research fields.

## Methods

### Sample collection, fixation, and resin embedding

*Arabidopsis thaliana* plants were cultivated in unsterilized soil [SupermixA (Sakata seed Co.) : vermiculite, 1:1] within 4-cm PolyPods. Two-week-old seedlings and soil were fixed in pre-fixative (4% formaldehyde, 2% glutaraldehyde in cacodylate or phosphate buffer) at 4 °C overnight after degassing. Rhizosphere soil from field beds at RIKEN Yokohama campus (collected May 2023, June 2023, April 2024) and a colony of moss (species not identified, collected May 2023) were collected and similarly fixed. Samples were post-fixed with 1% osmium tetroxide for 4 h, dehydrated through an ethanol series (25%, 50%, 75%, 90%, 100%), transitioned through propylene oxide to promote resin penetration, and embedded in a low viscosity epoxy resin (Quetol 651, Nissin EM) at 70 °C for 96 h.

### Sectioning and polishing

The embedded samples were sectioned into 3- or 4-mm slices using a diamond band saw (EXAKT 300, EXAKT Co.) and polished using an automatic mechanical polisher (MA-150, Musashino Electronics Co.). Sequential polishing was performed using P1000 and P4000 abrasive papers for 20 min each. As an optional polishing, flat milling was performed for 3 min using an ion milling machine (Hitachi ArBlade 5000).

### Electron staining and SEM observation

The polished blocks were stained with 0.4% uranyl acetate (10 min)^28^, rinsed, lead stained (2 min, lead stain solution, Sigma), and dried. Alternatives included 0.4% samarium chloride or Meyer’s hematoxylin (10 min, Merck), yielding comparable contrast³. The dried specimens were mounted with carbon paste and coated with osmium (HPC-1SW, Vaccum Device Co.). Imaging was performed using FE-SEM (SU8220 or SU8600, Hitachi High-Tech Co.) equipped with YAG-BSE (SU8220, 5 kV)^28,29^ or OCD-BSE (SU8600, 5–10 kV) LA-BSE (2 kV) detectors. The low-magnification surface quality of the resin blocks before and after polishing was examined using a tabletop SEM (TM4000, Hitachi High-Tech Co.) operated at 5–10 kV in the BSE low-vacuum mode.

The full protocols and comparative results are detailed in the Supplementary Information.

## Acknowledgements

We thank Dr. Ken Shirasu and the members of his laboratory (RIKEN CSRS) for their valuable advice on symbiosis. We also thank Dr. Hiroyuki Sasaki (RIKEN IMS) and Dr. Takeshi Matsui (Tokyo University of Technology) for their kind instruction for hematoxylin staining. We are grateful to all members of Meiwafosis Co., Ltd. and Musashino Denshi Inc. for their demonstrations of cutting and polishing, and to Hitachi High-Tech Co. for their support with SEM observation and ion milling.

## Authors’ Contributions

Y.S., Y.G., and M.S. prepared the samples and conducted the observations. K.T. designed and planned the experiments and observations, and wrote the manuscript.

