## Extended Data for "High-Resolution Imaging of the Rhizosphere Using Resin-Embedded Cross-Sectional Polishing and Scanning Electron Microscopy for Biological Samples (bioCP-SEM)"

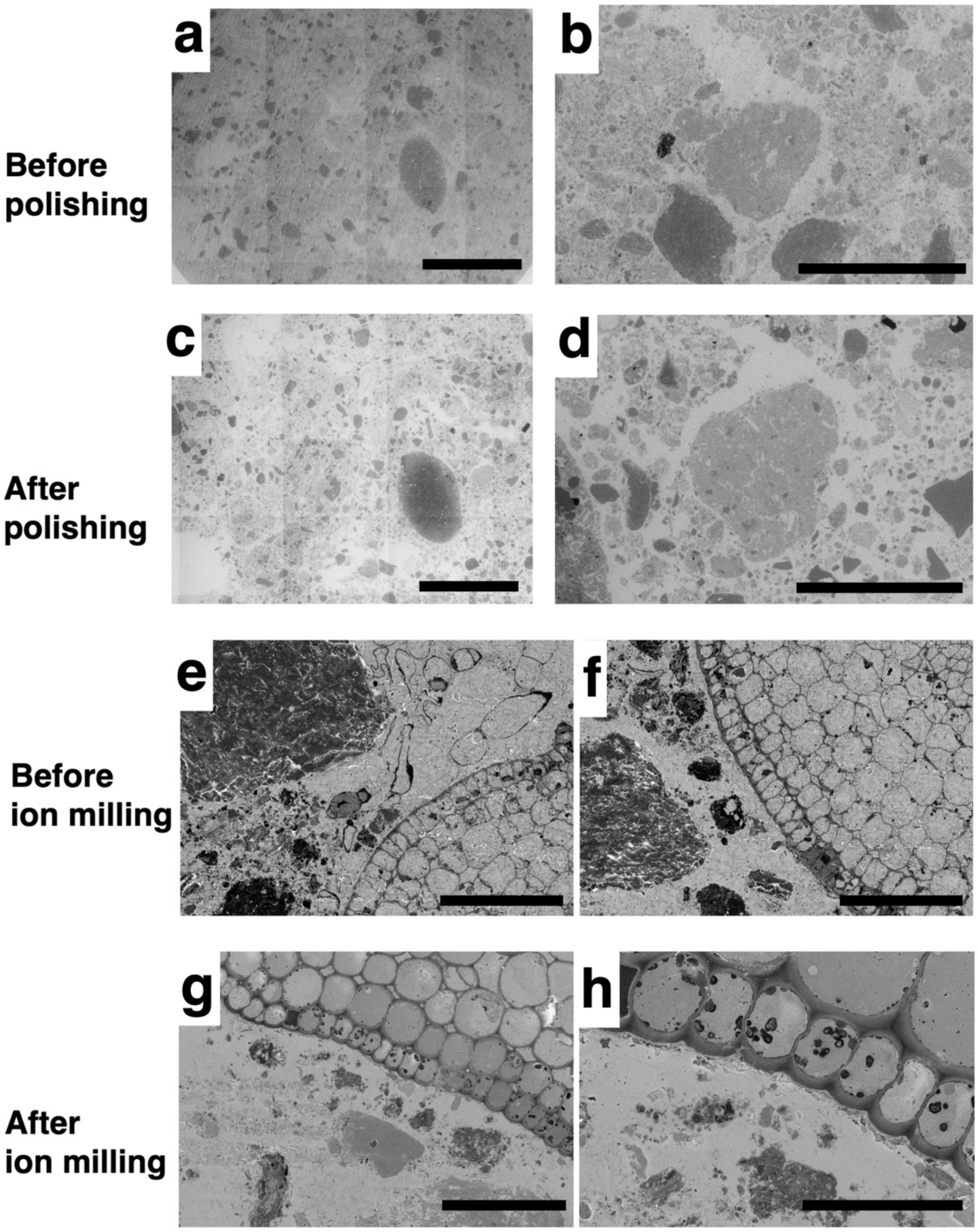


**Extended Data Fig. 1. Comparison of cross-sectional quality before and after polishing and ion milling.** (a–d) Comparison of surface quality before (a, b) and after (c, d) mechanical polishing. Polishing removed surface roughness and charging artifacts, resulting in improved contrast and visibility of fine structures. Images in (a–d) were acquired using a tabletop scanning electron microscope (Hitachi TM4000, 5 kV, BSE low vacuum mode). (e–h) Comparison of cross-sections before (e, f) and after (g, h) ion milling. Ion milling produced smoother sample surfaces, enabling clearer visualization of intracellular and tissue structures with reduced contamination and curtaining effects. Scale bars: (a, c) 2 mm; (b, d) 500 μm; (e–g) 100 μm; (h) 40 μm.

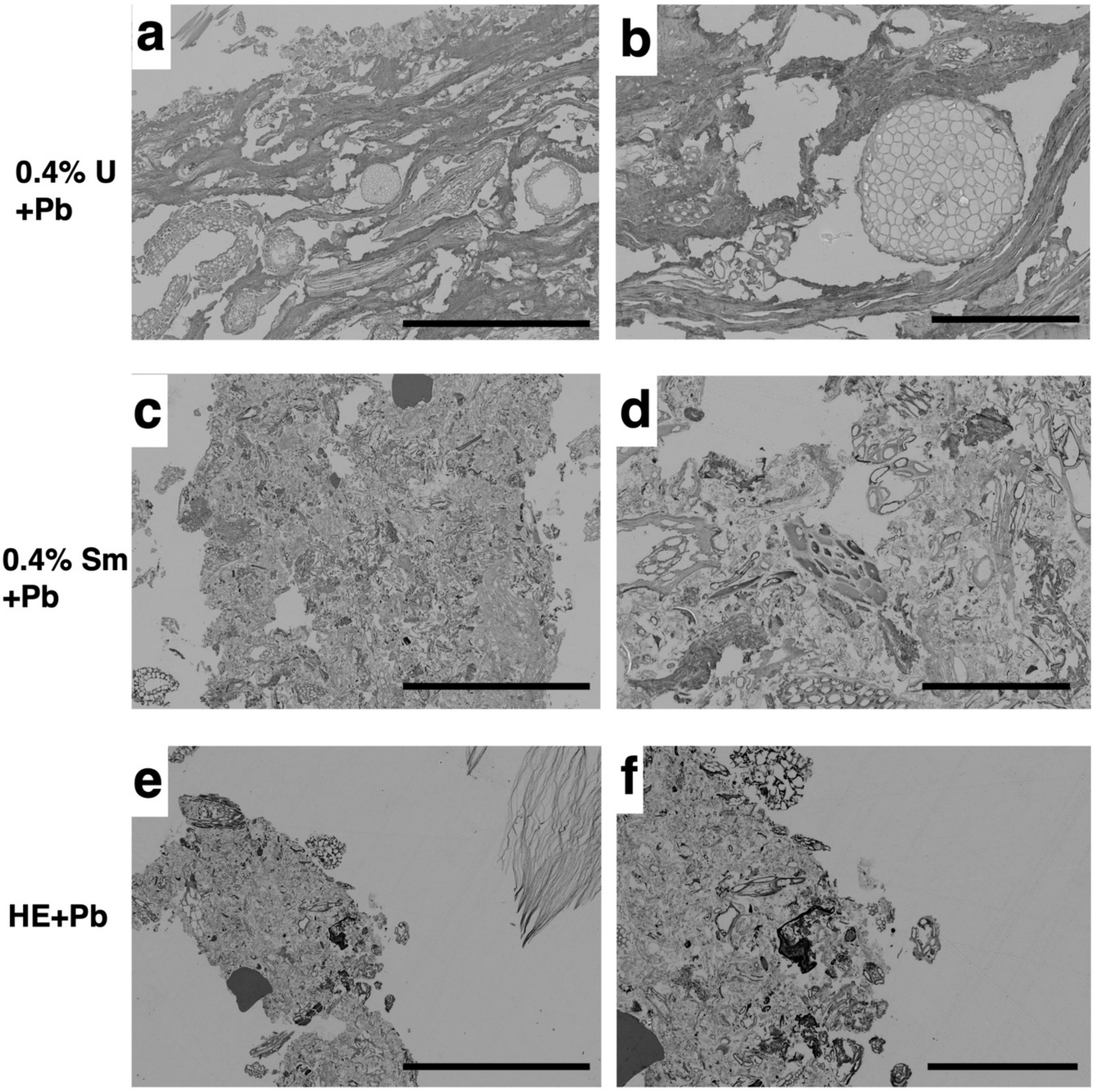


**Extended Data Fig. 2. Comparison of SEM image contrast in uranyl acetate, samarium chloride, and hematoxylin staining.** (a, b) Ultrastructural images stained with 0.4% uranyl acetate (U) and lead (Pb). (c, d) Ultrastructural images stained with 0.4% samarium chloride (Sm) and Pb. (e, f) Ultrastructural images stained with Mayer's hematoxylin (HE) and Pb. Scale bars: (a, c, e) 500 µm; (b, d) 100 µm; (f) 200 µm.
