## Supplementary Information for "High-Resolution Imaging of the Rhizosphere Using Resin-Embedded Cross-Sectional Polishing and Scanning Electron Microscopy for Biological Samples (bioCP-SEM)"

### 1. Sample fixation and resin embedding

#### Materials

- 4% paraformaldehyde and 2% glutaraldehyde in 0.05 M phosphate buffer or cacodylate buffer (pH 7.2)
- 1% osmium tetroxide in 0.05 M phosphate buffer or cacodylate buffer (pH 7.2)
- 0.05 M buffer (phosphate buffer or cacodylate buffer, pH 7.2)
- Ethanol series: 25%, 50%, 75%, 90%, 100% ethanol in distilled water
- Propylene oxide
- Epoxy resin: Quetol 651 (Nisshin EM)
- Polypropylene conical tubes (5–200 mL) with hole at the bottom or beakers

#### Protocol

1. Collect *Arabidopsis* plants (grown in 4-cm pots) or wild plant root–soil samples by coring or block cutting. Keep root–soil association intact.
2. Place samples into tubes or beakers with a small hole at the bottom to allow fluid exchange by gravity.
3. Slowly add fixative from the top (as if watering) to avoid disturbing the soil structure. Degas under vacuum (to -0.08~-0.1 MPa, gage pressure) for several short cycles, followed by 30–60 min under continuous vacuum to remove trapped air.
4. Fix samples overnight at 4 °C.
5. Wash six times with 0.05 M buffer (5 min each).
6. Post-fix in 1% osmium tetroxide for over 4 h at room temperature (22-24 °C).
7. Dehydrate samples in a graded ethanol series (25%, 50%, 75%, 90%, 100% EtOH, 3 × 30 min per step).
8. Substitution with propylene oxide:

   Propylene oxide:ethanol 1:1 (30 min), 100% Propylene oxide (2 × 1 h).

   Before infiltration, prepare Quetol 651 resin according to the manufacturer's formulation:

- Quetol 651 resin: 18.7 mL
- NSA (nonenyl succinic anhydride): 20.1 mL
- MNA (methyl nadic anhydride): 11.2 mL
- DMP-30 (2,4,6-tris(dimethylaminomethyl)phenol): 0.75 mL

  Mix thoroughly before use.

9. Infiltrate with low viscosity epoxy resin (e.g., Quetol651) gradually over 3 days:

   resin:propylene oxide 1:2 (12 h), 1:1 (12 h), 2:1 (12 h) → pure resin (3 × 12–24 h). After transitioning to 100% resin, degas again using several short cycles to -0.08~0.1 MPa followed by 30 min under vacuum to eliminate remaining trapped air and ensure complete infiltration.

10. Cure resin at 70 °C for over 96 h for Quetol651 to obtain solid resin blocks. For Quetol651, it is recommended to cover the resin surface with a sheet or film (e.g., polypropylene or polyethylene) to prevent the resin components from volatilizing during curing.

---

### 2. Sectioning and polishing

#### Equipment

- Diamond band saw (EXAKT 300, EXAKT Co.)
- Automatic polishing machine (MA-150, Musashino Electronics)
- P1000 and P4000 waterproof abrasive papers
- 1-μm diamond slurry
- Polishing cloth
- Ultrasonic cleaner
- Ion milling machine (ArBlade 5000, Hitachi)

#### Protocol

1. Secure cured resin blocks on the saw stage using jigs.
2. Cut blocks to 3–5-mm thickness using ±15° arc motion (“contact point method”).

Apply water to reduce heat and debris.

3. Mount cut surface downward onto polishing stage.
4. Polish using the following sequence:
    - P1000 abrasive paper for 10–20 min with water
    - P4000 paper for 10–20 min with water
    - Optional: 1-µm diamond slurry on cloth for 20 min or flat-milling for 3 min using an ion milling machine (Hitachi ArBlade 5000). Rinse sections in distilled water and ultrasonicate for 3 min.
5. Rinse sections in distilled water.

---

#### 3. Electron staining and conductive coating

##### Staining options

- **Standard**:
  0.4% uranyl acetate (10 min) → wash (DW)→ lead stain solution (2 min) → wash (DW)
- **Non-radioactive alternatives**:

0.4% samarium chloride (10 min) → wash (DW) → lead stain solution (2 min) → wash (DW)

Meyer's hematoxylin (20 min, filtered through a 0.45-µm Millipore filter) → wash (DW) → lead stain solution (2 min) → wash (DW)

**Note**: For large area sections, non-radioactive options are preferable due to uranyl usage limitations.

##### Conductive coating

1. After staining, completely air-dry the polished sections.
2. Mount on aluminum stubs using carbon tape or carbon paste.
3. Coat with osmium (2–3 nm) using osmium plasma coater (HPC-1SW or equivalent).

---

### 4. SEM imaging and image processing

#### Equipment

- Field emission SEM (SU8220, SU8600; Hitachi High-Tech)
- Benchtop SEM (TM4000plusII; Hitachi High-Tech)
- Detectors: LA-BSE, YAG-BSE, OCD-BSE

#### Imaging conditions

| Detector | Accelerating voltage | Notes |
|---|---|---|
| YAG-BSE | 5–10 kV | High contrast for soft–hard interfaces |
| LA-BSE | 2–5 kV | Low-angle detail, good for surface |
| OCD-BSE | 10 kV | Higher resolution, newer systems |
| TM4000 BSE-L | 5 kV | For quick overview imaging |

#### Image adjustment

- Backscattered electron images appear as negative contrast compared to TEM (heavy = white).
- Use grayscale inversion in ImageJ or SEM software for TEM-like visualization.
- Contrast enhancement and scale bar calibration are performed before the final export.

---

#### Supplementary notes

- Sections thinner than 2 mm are prone to warping and are not recommended for this method.
- Microbial structures, such as coccoid clusters, hyphae, and arbuscules, and EPS-rich biofilms can be visualized at submicron resolution.

#### Figure for supplementary information

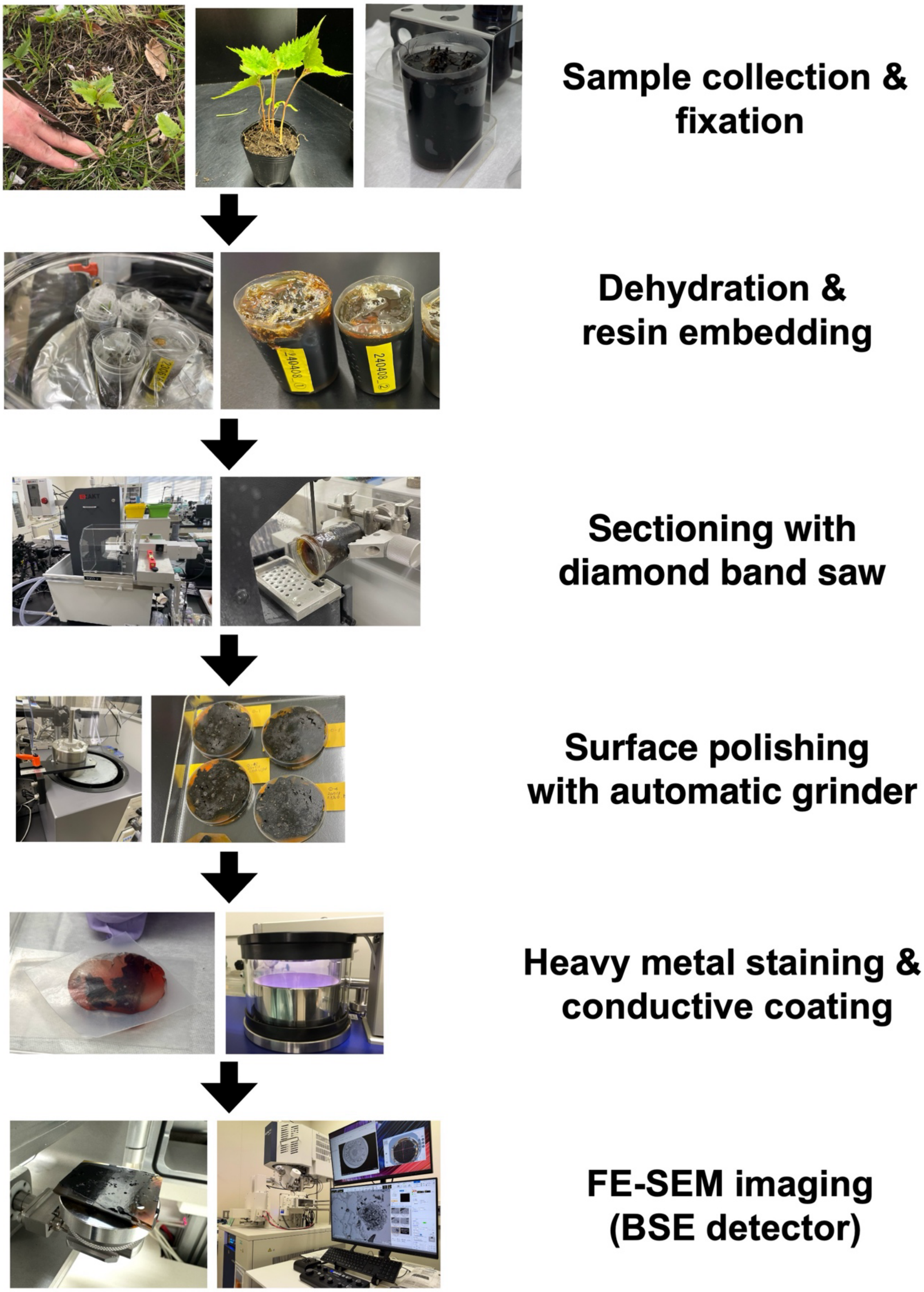


**Workflow of the bioCP-SEM method.** Sequential steps for preparing soil–plant blocks for backscattered electron (BSE) imaging by field-emission scanning electron microscopy (FE-SEM). Samples (plants with surrounding soil) were collected and chemically fixed

(Sampling & fixation), followed by dehydration and embedding in epoxy resin (Dehydration & embedding in resin). Resin-embedded blocks were cut into appropriate sizes using a diamond band saw (cutting with diamond band saw) and subsequently polished to obtain a flat cross-section using an automatic grinder (polishing with auto grinder). After surface metal staining and conductive coating (metal staining and coating), the samples were mounted onto specimen holders and imaged using FE-SEM equipped with a BSE detector (FE-SEM observation). This workflow enables high-resolution visualization of roots, microbes, and minerals in situ within the rhizosphere.
